# Rational selection of TbpB variants elucidates a bivalent vaccine formulation with broad spectrum coverage against *Neisseria gonorrhoeae*

**DOI:** 10.1101/2024.09.07.611798

**Authors:** Jamie E. Fegan, Epshita A. Islam, David M. Curran, Dixon Ng, Natalie Au, Elissa G. Currie, Joseph Zeppa, Jessica Lam, Anthony B. Schryvers, Trevor F. Moraes, Scott D. Gray-Owen

**Author notes:** These authors contributed equally to this work (Jamie E. Fegan, Epshita A. Islam). Correspondance (ABS); (TFM); or (SDG).

## Abstract

*Neisseria gonorrhoeae* is the causative agent of gonorrhea, an on-going public health problem due in part to the lack of success with efforts to develop an efficacious vaccine to prevent this sexually transmitted infection. An attractive candidate vaccine antigen because of its essential function and surface exposure, the gonococcal transferrin binding protein B (TbpB) exhibits high levels of antigenic variability which poses a significant obstacle in evoking a broadly protective vaccine composition. Here, we utilize phylogenetic information to rationally select TbpB variants for inclusion into a potential gonococcal vaccine and identify two TbpB variants that when formulated together elicit a highly cross-reactive antibody response in both rabbits and mice against a diverse panel of TbpB variants and clinically relevant gonococcal strains. Further, this formulation performed well in experimental proxies of real-world usage, including eliciting bactericidal activity against 8 diverse gonococcal strains and decreasing the median duration of colonization after vaginal infection in female mice by two heterologous strains of *N. gonorrhoeae*. Together, these data support the use of a combination of TbpB variants for a broadly protective gonococcal vaccine.

## Introduction

*Neisseria gonorrhoeae* (Ngo) causes the sexually transmitted infection gonorrhea, which has been successfully treated with antibiotics for decades. Worryingly, an increasing prevalence of antibiotic resistant Ngo threatens our ability to successfully treat this infection^1^, leading to the potential of untreatable disease by multi-drug resistant strains^2,3^. This has created a strong impetus to develop new countermeasures to combat gonococcal infections. The modest but promising cross-protection against gonococcal infection elicited by the MeNZB outer membrane vesicle vaccine^4^ and the multicomponent meningococcal B vaccine 4CMenB (Bexsero)^5–7^ has demonstrated that vaccine-mediated protection against the gonococcus is feasible, and has reinvigorated efforts to develop an efficacious, broadly protective vaccine.

As the gonococcus lacks a polysaccharide capsule, most vaccine efforts have focused on surface-exposed proteins, including gonococcal pili^8^, which failed to elicit protection in male volunteers. One long-standing candidate is the bacterial transferrin receptor, composed of the integral outer membrane protein transferrin binding protein A (TbpA) and the surface-anchored lipoprotein transferrin binding protein B (TbpB). TbpA is vital for the gonococcus to acquire the essential micronutrient iron from the host protein transferrin (Tf)^9^, while TbpB specifically binds iron-loaded (holo) human Tf (hTf) to increase the efficiency of iron uptake^10^. The gonococcal transferrin receptor was shown to be essential for male urethral infection in humans by strains of Ngo that lack a lactoferrin receptor^11,12^, thus both transferrin binding proteins have been posited to be attractive candidate vaccine targets. While TbpA exhibits far less sequence variation compared to TbpB^13^, inherent difficulties exist in targeting integral membrane proteins, including protein stability, large scale protein production, and methods of focusing the immune response to specifically target surface exposed regions of the protein. In contrast, TbpB is a soluble, stable lipoprotein, making commercial production and formulation of a TbpB-based vaccine more tractable. However, TbpB exhibits substantial antigenic variation^14,15^, presumably due to its accessibility to the immune system since its lipidated and flexible peptide anchor allows it to extend away from the bacterial surface to capture transferrin.

TbpB from the closely related pathogen *Neisseria meningitidis* (Nme) has been extensively evaluated for the diversity of variant sequences and cross-reactivity of antibodies elicited after immunization^16^. Meningococcal TbpBs fall into two distinct phylogenetic clusters exhibiting low sequence identity and eliciting minimal cross-reactivity; the smaller isotype I TbpBs (1.8 kb gene, approximately 68 kDa protein) and the larger and more diverse isotype II TbpBs (2.1 kb gene, approximately 80-90 kDa protein)^17^. A crystal structure of a TbpB from each meningococcal cluster has been solved (isotype I, strain B16B6, PDB ID #4QQ1^13^; isotype II, strain M982, PDB ID #3VE2^18^), both of which show a bi-lobed protein containing an N and C lobe, with each lobe consisting of a handle and barrel domain. The N lobe of TbpB is responsible for binding to the C lobe of iron-loaded hTf^19,20^ and is more variable than the well conserved C lobe^13^. Gonococcal TbpB variants all fall within the meningococcal isotype II TbpB cluster, forming two distinct subclusters^21^. Thus, while there is still antigenic variability to contend with, it is expected that this variability is manageable with respect to the selection of variants for inclusion into a vaccine formulation.

Here, we evaluate the potential of combining multiple gonococcal TbpB variants to elicit broad cross-reactivity. We use a phylogenetically informed selection of variants to generate vaccine formulations consisting of different combinations of TbpBs, quantify the cross-reactive and bactericidal propoerties of the elicited antibodies, and demonstrate protection against gonococcal challenge in the murine lower genital tract colonization model. Combined, this work reinforces the potential to develop a broadly protective TbpB-based gonococcal vaccine.

## Results

### Unbiased selection of TbpB antigens based on phylogentic analysis

Gonococcal TbpBs exhibit antigenic variation that separates into two distinct phylogenetic clusters (Fig 1A). The larger of these, which we have termed Ngo cluster 1, is most similar to the previously defined Nme cluster 3, which contains the TbpBs from meningococcal strains H44/76 and N224, and the smaller gonococcal cluster which we have termed Ngo cluster 2, which is most similar to TbpBs from Nme cluster 2, containing TbpBs from meningococcal strains MC58 and M982 (Fig 1B).

**Figure 1:**
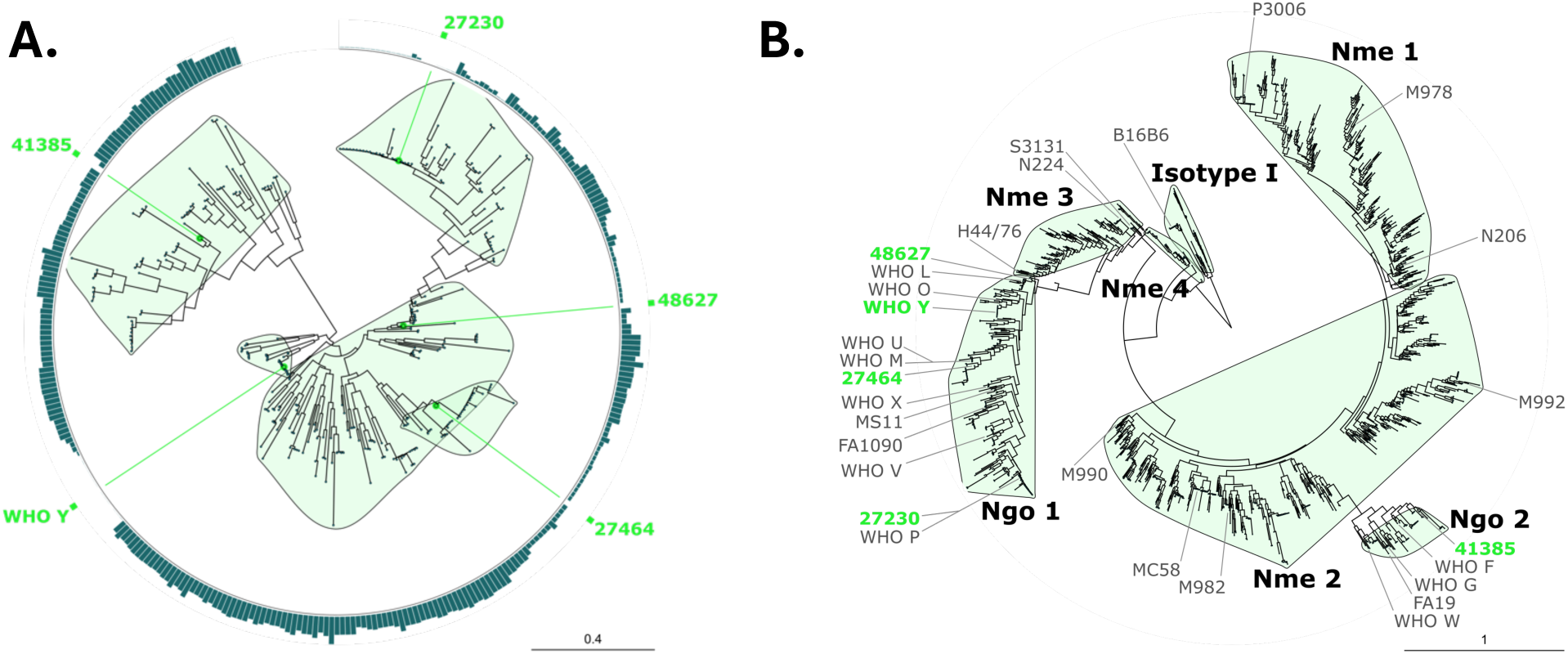
Bioinformatic selection of representative TbpBs. **A.** Phylogenetic tree of 325 Ngo TbpB sequences, clustered into 5 clades. The representative sequence selected for each clade is indicated in green. Gonococcal cluster 1 variants are on the right and encompass the 27230, 48627, 27464, and WHO Y clades, while the gonococcal cluster 2 variants consist of the 41385 clade in the upper left. **B.** Phylogenetic tree of 1485 TbpB sequences from Ngo and Nme, clustered into 7 descriptive clades. The 2 gonococcal clades are found on opposite sides of the tree, and are highly dissimilar. The Nme sequences are clustered into 5 clades, with the Isotype I sequences as the most dissimilar to anything else in the tree. The Nme 3 and Nme 4 clades are well separated and distinct, while the large Nme 1 and Nme 2 clades contain more of a gradient of sequence groups. The 5 representative Ngo sequences selected for further characterization are indicated in green, and the 24 sequences indicated in grey were utilized in the cross-reactivity panel in figures 2, 3, 4, and 6.

As a first step towards evaluating potential gonococcal TbpB variants for their ability to elicit the broad cross-reactivity, we selected 5 representative Ngo TbpB sequences using the phylogenetic analysis software Navargator (Fig 1A)^22^. This program takes a phylogenetic tree as input and selects a designated number of representative sequences such that they are as close as possible to as many other sequences in the phylogenetic cluster they represent. Selected variants are summarized in Table 1 and broken down by which of the two gonococcal clusters they belong to.

**Table 1:**
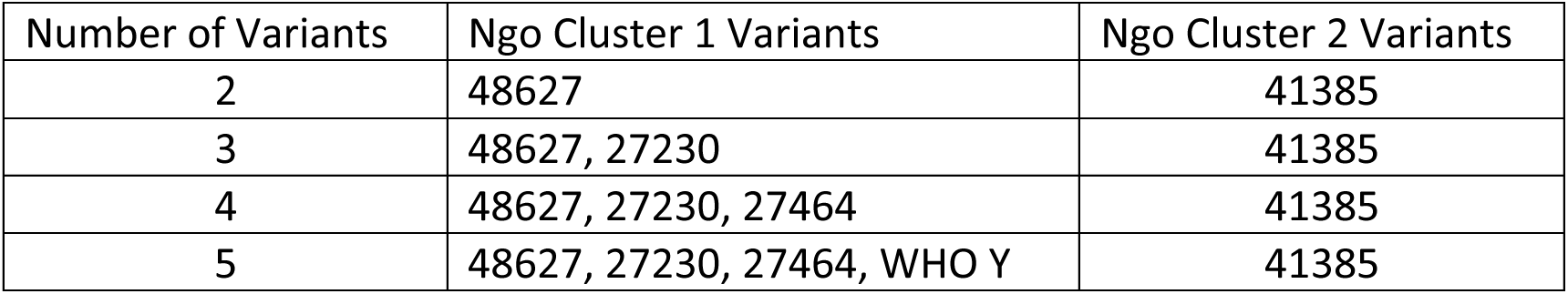
Variant selection for gonococcal TbpBs based on the number of requested variants.

With this, we considered what impact the cumulative addition of variants would have on cross-reactivity of the antibody response against the known diversity of TbpB variants. The selected *tbpB* genes were codon optimized for expression in *E. coli*, synthesized, and then recombinantly expressed and purified from *E. coli*. Individual TbpBs, a bivalent composition consisting of a single representative TbpB from each of the two major gonococcal clusters (Ngo TbpBs 48627 and 41385), and a pentavalent composition that included all five representative TbpBs were formulated with aluminum hydroxide gel. Details of the formulations given to each rabbit, including antigen amount, can be found in Table 2. These were sub-cutaneously injected into rabbits three times (day 0, 21, and 42) and blood samples were acquired prior to each dose of vaccine and then again as a terminal bleed two weeks after the third dose. These were subsequently used to assess antiserum cross-reactivity against TbpB variants from both Ngo and Nme.

**Table 2:** Vaccine compositions for New Zealand rabbits.

| Rabbit | TbpB variant | Amount of protein dose 1 (per antigen) | Total protein dose 1 | Amount of protein doses 2 and 3 (per antigen) | Total protein doses 2 and 3 | Adjuvant |
| --- | --- | --- | --- | --- | --- | --- |
| 1 | 48627 | 100 μg | 100 μg | 50 μg | 50 μg | AlOH* |
| 2 | 41385 | 100 μg | 100 μg | 50 μg | 50 μg | AlOH |
| 3 | 27464 | 100 μg | 100 μg | 50 μg | 50 μg | AlOH |
| 4 | 27230 | 100 μg | 100 μg | 50 μg | 50 μg | AlOH |
| 5 | WHO Y | 100 μg | 100 μg | 50 μg | 50 μg | AlOH |
| 6 | 48627 + 41385 | 100 μg | 200 μg | 50 μg | 100 μg | AlOH |
| 7 | All 5 | 50 μg | 250 μg | 25 μg | 125 μg | AlOH |
| 8 | 48627 | 100 μg | 100 μg | 50 μg | 50 μg | Addavax |
| 9 | 41385 | 100 μg | 100 μg | 50 μg | 50 μg | Addavax |

\*, Aluminum hydroxide gel

### Formulations with two or more TbpBs elicit broadly cross-reactive antibodies with transferrin blocking activity

Pre-immune serum along with serum collected post 1, 2, and 3 doses of vaccine were assessed from each rabbit using high throughput protein ELISAs^23^. To represent the natural diversity of neisserial TbpBs, the ELISA panel consisted of 18 recombinantly-expressed gonococcal TbpBs split between Ngo cluster 1 and Ngo cluster 2 (n=13 representatives for Ngo cluster 1, n=5 representatives for Ngo cluster 2), and 11 meningococcal TbpB variants (n=1 isotype I TbpB, n=10 isotype II TbpBs split across the four isotype II clusters (n=3 for Nme cluster 1, n=4 for Nme cluster 2, n=3 for Nme cluster 3)) (Figs 1B and 2A). Binding of HRP-labelled hTf (hTf-HRP) was used as an expression control, since TbpBs binding ability to hTf-HRP indicates that they were present and properly folded on the ELISA plate (Fig 2B). As expected, pre-immune serum (0 doses) elicited minimal reactivity to any TbpBs included in the panel, while the breadth and intensity of reactivity generally increased with each dose (Fig 2A). Serum from rabbits that received a single antigen formulation had reactivity against a a subset of the gonococcal TbpBs, with the formulation containing Ngo TbpB 48627, the main representative of the largest TbpB cluster (Ngo cluster 1), having the broadest reactivity after three doses. None of the single TbpB formulations elicited reactivity that extended to broadly recognize the meningococcal TbpBs. In contrast, formulations composed of either the bivalent TbpB formulation (Ngo 48627 + Ngo 41385) or the pentavalent TbpB formulation containing all five TbpBs were broadly reactive against the full gonococcal TbpB panel and, notably, extended the anti-TbpB cross-reactivity to include the meningococcal TbpBs. Despite containing three additional TbpB variants beyond the bivalent formulation, the pentavalent formulation did not substantially alter the breadth of cross-reactivity, particularly when focusing on reactivity with the Ngo TbpBs. Interestingly, there was minimal change in the reactivity of sera from post-two doses as compared to sera from post-three doses, indicating that there may be limited maturation of the antibody repertoire between the second and third dose in the time span evaluated.

**Figure 2:**
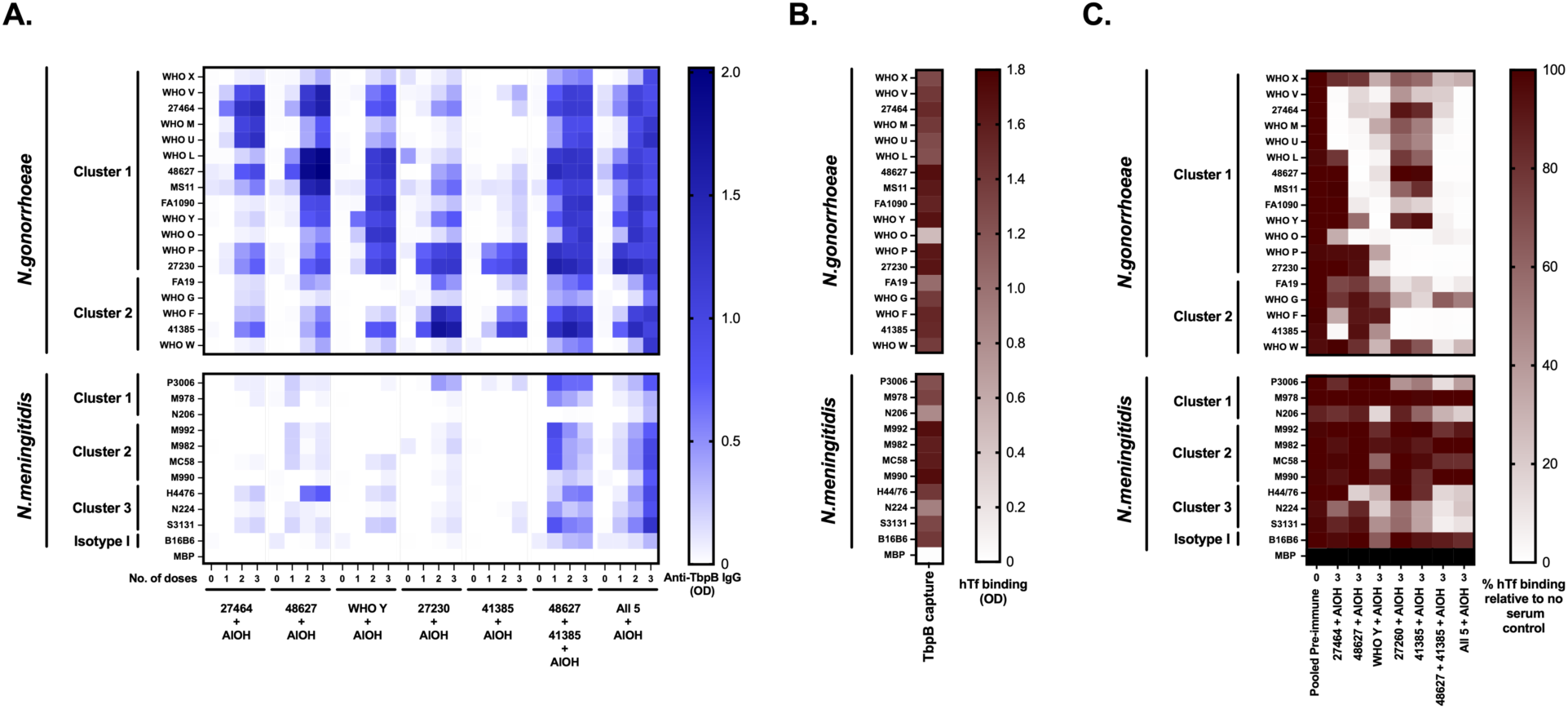
Coverage of pan-Neisserial TbpB diversity using gonococcal TbpBs as vaccine antigens. **A.** Heat map depicting ELISA-based reactivity of rabbit antiserum against a diverse panel of Neisserial TbpBs. Each row includes a different TbpB protein as capture antigen, with the phylogenetic cluster indicated to the left. Each column represents either pre-immune or post 1, 2 or 3 vaccine dose serum for a single rabbit immunized with the vaccine formulation indicated below the x-axis. Increasing intensity indicates increasing immunoglobulin G specific for that TbpB variant. Background noise from empty vector control (maltose binding protein, MBP) has been subtracted. **B.** hTf-binding to each TbpB as a plate control to quantification of properly folded (functional) TbpB. **C.** Heat map depicting the ability of post 3 dose immune serum to block hTf binding to the TbpB capture antigen. Values are normalized to the signal obtained from no serum control. Each column represents a different rabbit immunized with the vaccine indicated below, or pooled pre-immune serum as a baseline control. Black represents lack of hTf binding for the MBP empty vector control. AlOH, Aluminum hydroxide. Heat maps were generated using GraphPad Prism 10.1.1.

Aside from the significance of inducing broadly cross-reactive antibodies through TbpB immunization, it is postulated that the ability of antibodies to block the interaction between TbpB and hTf may lead to a reduced ability of the gonococcus to acquire iron. Consequently, this could contribute to protection via nutritional immunity, wherein the bacteria are starved of iron^24,25^. Therefore, we next assessed whether serum antibodies were able to inhibit HRP-conjugated hTf binding to TbpBs on the ELISA plate. Notably, as these immunizations were done in rabbits and rabbit Tf is not able to bind to Ngo TbpBs^26^, endogenous rabbit Tf is not expected to interfere with antibody development during immunization or to interfere during in vitro analysis. Since the quantity of pre-immune serum was limited, we utilized pooled pre-immune sera as the negative control and observed no reduction in HRP-hTf signal, as expected (Fig 2C). In contrast, terminal serum led to a substantial reduction in HRP-hTf binding, as denoted by lighter colour on the heat map. Single TbpB formulations elicited variable levels of hTf blocking against the gonococcal TbpBs, with Ngo WHO-Y eliciting the broadest hTf blocking among the single formulations, and Ngo 48627 eliciting the next best blocking ability. Among all the single TbpB formulations, blocking was highest against TbpBs that had the highest antibody reactivity, indicating that higher antibody binding leads to increased hTf blocking, as expected. Both the bivalent and the pentavalent TbpB formulations elicited very broad hTf blocking, and this again extended activity into a subset of the meningococcal TbpBs, which was not generally seen with single TbpB formulations. Again, consistent with the TbpB cross-reactivity, there was no substantial difference in TbpB-hTf blocking activity between the bivalent and pentavalent TbpB formulations.

### Alum and emulsion based formulations elicit similar cross-reactivity and transferrin blocking pattern

As the bivalent composition containing Ngo 48627 and 41385 TbpBs elicited broad coverage against the panel of TbpBs, we next evaluated the effect of adjuvant on the breadth of reactivity on these two antigens as single formulations. While the alum-based Alhydrogel was used for our initial evaluation, we additionally tested the oil-in-water emulsion-based adjuvant Addavax (research grade MF59) to consider whether the coverage we observed remains consistent with a different choice of adjuvant. Serum from rabbits immunized with either Ngo 48627 or 41385 TbpB formulated with Addavax were compared to serum from the rabbits immunized with the same TbpBs formulated with Alhydrogel using the same assays as Figure 2. As can be seen in Figure 3A, the choice of adjuvant in the formulation did not substantially impact the breadth or pattern of reactivity, though reactivity after one dose of vaccine was slightly higher in sera from the rabbits that received the formulations containing Addavax. Similarly, the adjuvant did not substantially influence hTf blocking activity of the antiserum, and still only elicited blocking against gonococcal TbpBs (Fig 3B). The hTf blocking activity was again higher against the cluster that the TbpB originated from, as expected, and was primarily seen after two and three doses of vaccine.

**Figure 3:**
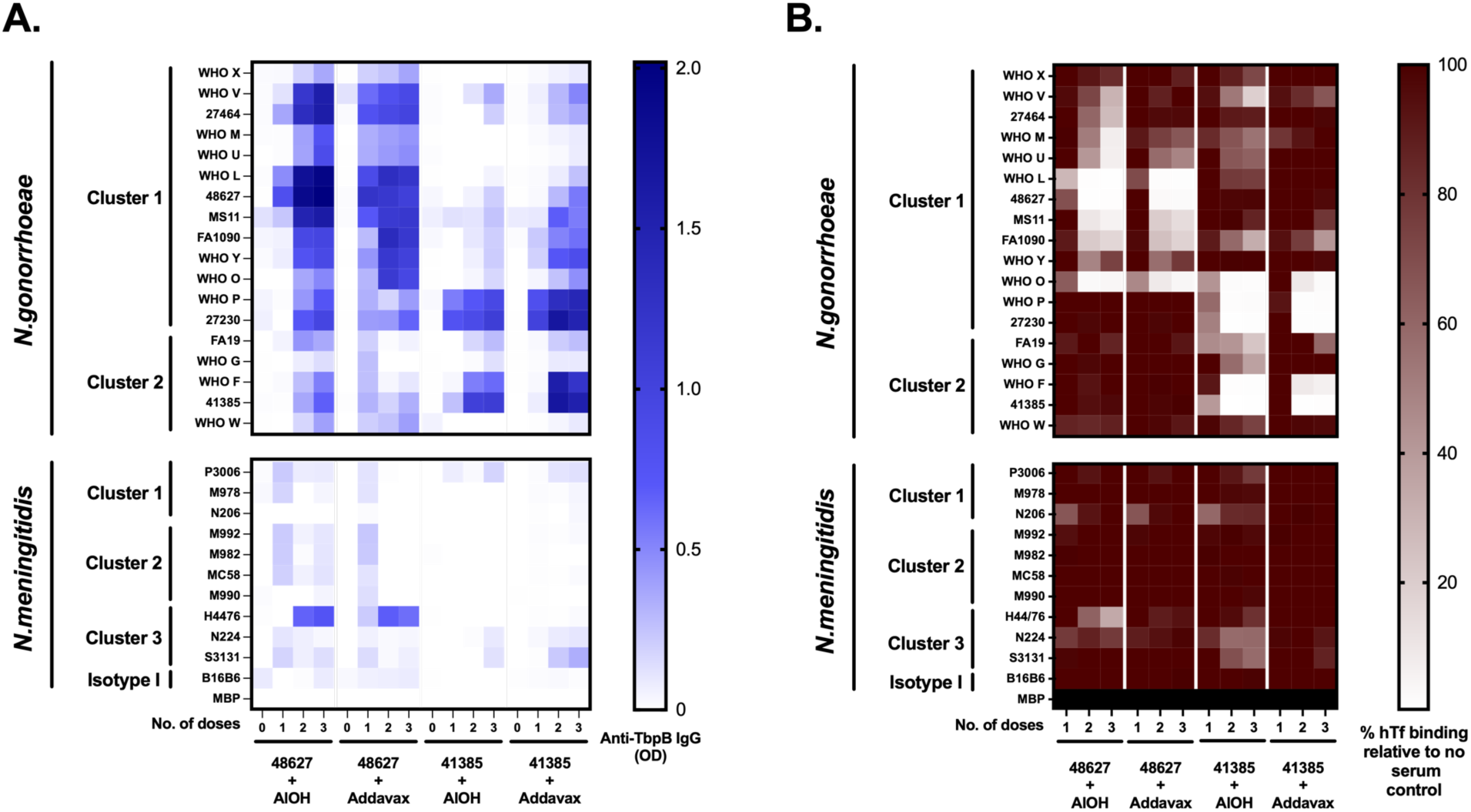
Coverage of TbpB diversity using antigens formulated with alum versus emulsion adjuvants. **A.** Heat map depicting reactivity of rabbit antiserum against a broad ELISA panel of Neisserial TbpBs. Each row includes a different TbpB capture antigen, with the phylogenetic cluster indicated to the left. Each column represents either pre-immune or post 1, 2, or 3 dose serum for a single rabbit immunized with the vaccine indicated below. Increasing intensity indicates increasing immunoglobulin G specific for that TbpB variant. Background noise from empty vector (MBP) has been subtracted. **B.** Heat map depicting the ability of post 1, 2, and 3 dose immune serum to block hTf binding by the TbpB capture antigen. Values are normalized to the signal obtained from no serum control. Each column represents a different rabbit immunized with the vaccine formulation indicated below. Black represents lack of hTf binding for the MBP empty vector control. AlOH, aluminum hydroxide. Heat maps were generated using GraphPad Prism 10.1.1.

### TbpB-based compositions provide broad recognition of clinically relevant gonococcal isolates

To consider the ability of a TbpB-based vaccine to elicit antibodies that recognize the gonococcal surface, a whole cell ELISA panel was developed to assess serum reactivity against 36 gonococcal strains, including three commonly used lab strains (MS11, FA1090, and FA19) and the WHO reference panel strains^27^, which contain some of the TbpB variants assessed in Figures 2 and 3, along with disease isolates acquired from Public Health Ontario (Canada) and a collection of gonococcal strains acquired from endocervical swabs from female sex workers in Nairobi, Kenya (Kenyan Ngo Collection)^28^. Additional information regarding the disease isolates can be found in supplemental Table 2. Included in the Public Health Ontario strains are recent isolates that lead to mucosal infections as well as disseminated gonococcal infections. Strains were grown under iron restricted conditions via chelation by deferoxamine to induce expression of the transferrin receptor and then heat killed prior to adherence to ELISA plates via drying. Binding of HRP-hTf was used as a control for receptor expression (Fig 4B), although this control cannot differentiate between TbpA and TbpB expression based upon hTf binding alone. More variability in hTf binding is apparent here compared to what was seen with the recombinantly expressed TbpB proteins in Figures 2 and 3. This may reflect inherent differences in expression and/or different levels of surface transferrin receptor expression in the culture conditions employed to grow the bacteria used to coat the ELISA wells. Regardless, all strains showed some level of hTf binding (Fig 4B), and all showed some reactivity with the TbpB-specific antisera (Fig 4A).

**Figure 4:**
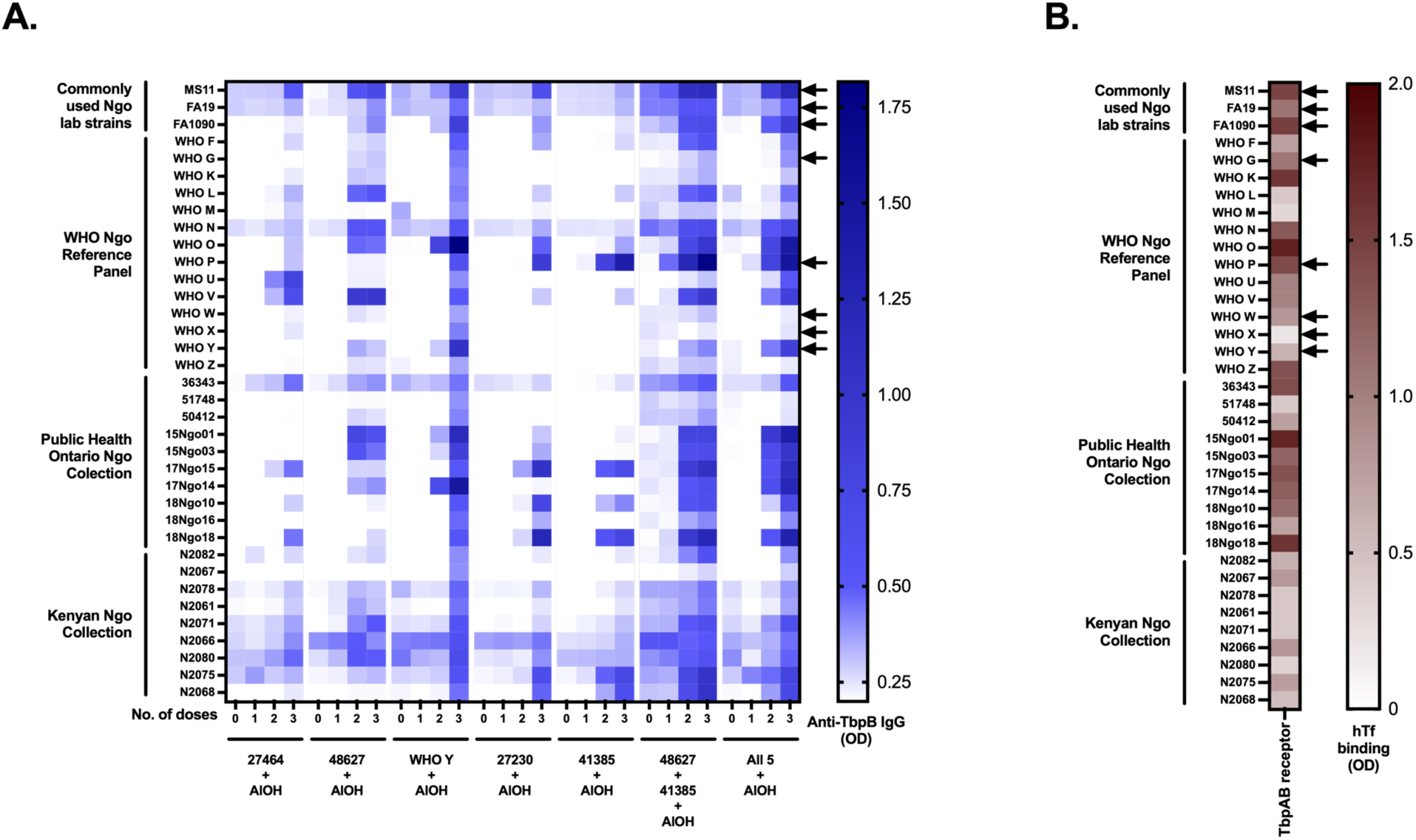
Coverage of clinical gonococcal isolates using gonococcal TbpBs as vaccine antigens. **A.** Heat map depicting reactivity of rabbit antiserum against a broad ELISA panel consisting of heat inactivated whole bacteria. Each row is a different gonococcal isolate, with the originating collection indicated to the left. Each column represents either pre-immune, or post 1, 2, or 3 dose serum for a single rabbit immunized with the vaccine indicated below. AlOH, aluminum hydroxide. **B.** hTf-binding to each isolate used as a readout for surface expression of the TbpAB receptor. Heat maps were generated using GraphPad Prism 10.1.1. Arrows indicate strains that were used for serum bactericidal assays shown in Figure 5.

Whole cell reactivity was variable based on the immunizing TbpB, with serum raised against Ngo WHO Y TbpB showing the broadest reactivity of the single protein formulations, while reactivity with either the bivalent or pentavalent TbpB formulations showed broad reactivity after only two doses of vaccine (Fig 4A). Interestingly, there was high reactivity by the bivalent formulation against the clinical isolates (Public Health Ontario and Kenyan Collections) that was not improved by the inclusion of the additional three TbpBs present in the pentavalent formulation.

### TbpB-based compositions elicit bactericidal antibodies

Eight gonococcal strains, consisting of three common laboratory strains (MS11, FA1090, and FA19) and five WHO strains (WHO G, P, W, X, and Y), were chosen to assess the bactericidal activity of immune rabbit serum (Fig 5). These strains were chosen to include representatives from each gonococcal TbpB cluster, porin type (which may influence gonococcal susceptibility to the bactericidal activity of human serum^29^), as well as to account for the varying degree of antibody reactivity and hTf binding (as denoted by arrows in Figure 4). Normal human serum (NHS) without antibody depletion was utilized as the source of complement in the bactericidal assays. Prior to the conducting these assays, NHS was titrated (0.5% to 15%) against individual strains to determine the highest percentage of NHS that did not impede Ngo recovery in the absence of an exogenous antibody source. Thus, bactericidal assays were conducted on strains that are highly resistant (FA1090 and WHO G, 15% NHS), intermediate resistant (WHO P, 5% NHS), and sensitive (MS11, FA19, WHO W, WHO X, and WHO Y, 2.5% NHS) to NHS.

**Figure 5:**
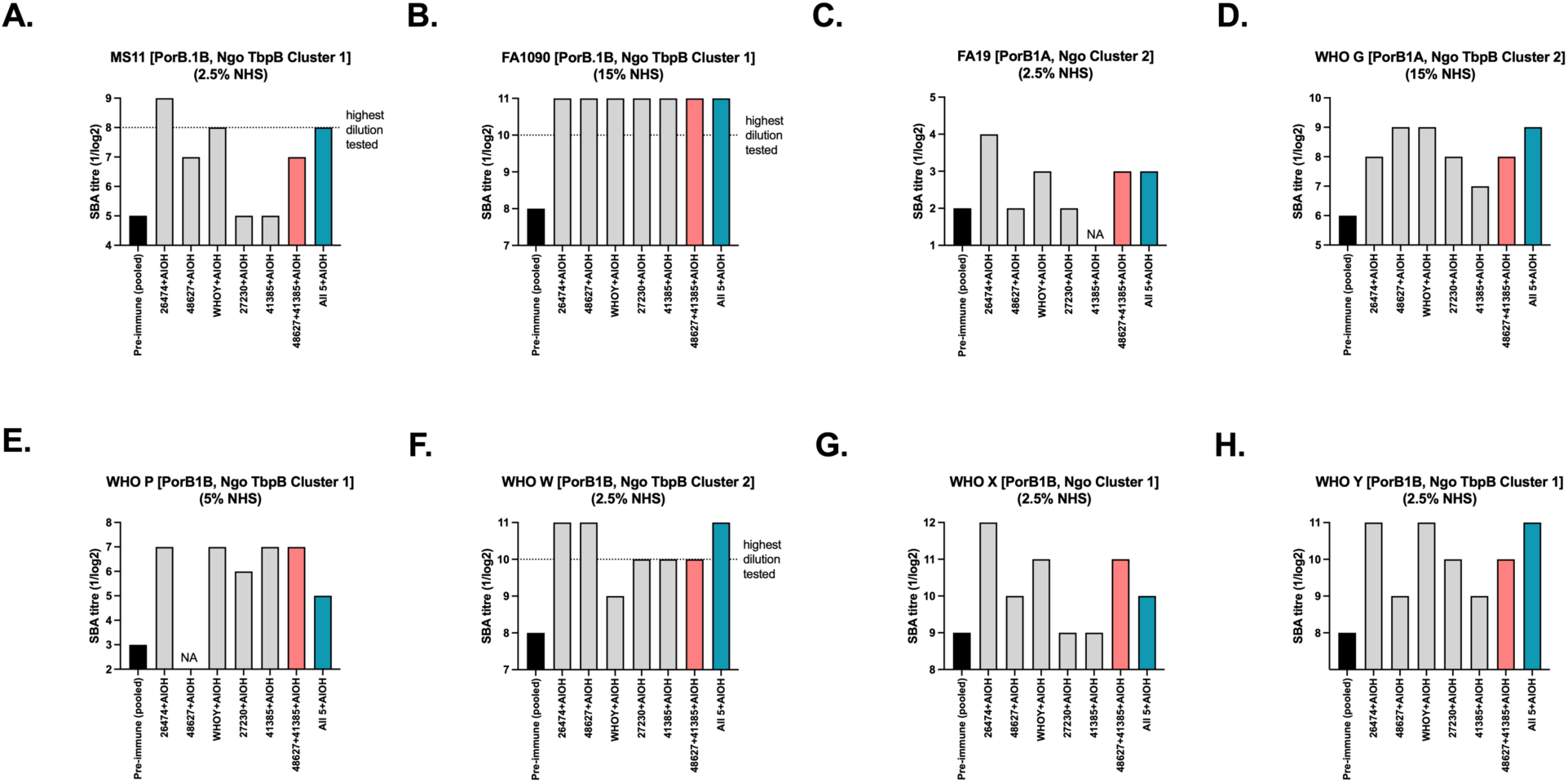
Serum bactericidal activity of post 3 dose rabbit serum against gonococcal isolates. **A-H.** Different *N. gonorrhoeae* strains were tested for bactericidal killing with the porin type, TbpB cluster, and percentage of normal human serum (NHS) used in the assay indicated above each panel. Bars represent serum bactericidal activity (SBA) Log2 titre (50% killing compared to no antibody control) for pooled pre-immune serum (black), post dose 3 serum from rabbits immunized with a single (grey), bivalent (pink), or pentavalent (teal) TbpB variant composition. Dotted line indicates highest dilution tested, where bars that exceed that line indicate that less than 50% killing was not obtained at any serum dilution tested.

High levels of bactericidal activity were seen in sera from rabbits immunized with any one of the single TbpB formulations when tested against Ngo FA1090, WHO G, WHO W, and WHO Y (Fig 5B, D, F, and H). The remaining strains (MS11, FA19, WHO P, and WHO X) were killed by some single TbpB-based formulations but not others (Fig 5A, C, E, and G). Notably, sera from bivalent and pentavalent TbpB formulations elicited bactericidal activity against all eight strains. The bivalent TbpB formulation elicited bactericidal activity one dilution lower than the pentavalent formulation for four strains tested (MS11, WHO G, WHO W, and WHO Y), equally to the pentavalent formulation for two strains tested (FA1090 and FA19), and better compared to the pentavalent formulation for the final two strains tested (WHO P and WHO X). These data again indicate that a bivalent TbpB formulation is sufficient to elicit broad bactericidal antibody coverage against diverse gonococcal strains. Also notable is that strains WHO W and WHO X had minimal binding to rabbit serum in the whole cell ELISAs but were still effectively killed in the bactericidal assays, indicating that a combination of reactivity and functional assays are needed to fully understand potential effector functions of antibodies elicited by different vaccine formulations.

### Bivalent TbpB composition accelerates clearance of heterologous gonococcal strains from the lower genital tract of female mice

As the bivalent TbpB formulation was sufficient to elicit broad cross-reactivity against the full panel of TbpBs, against the panel of gonococcal strains, and at eliciting bactericidal antibodies that recognized and killed all eight gonococcal strains tested, we selected this composition to evaluate protection in the female genital tract colonization model in mice. Ngo 48627 and Ngo 41385 TbpBs were formulated together with Addavax, the emulsion-based adjuvant. Female C57Bl/6 mice were immunized three times intra-peritoneally with the bivalent formulation or with Addavax alone and serum samples were collected two weeks after each dose. Immunogenicity for each mouse was tested against the two component TbpBs, Ngo 41385 (Fig 6A) and Ngo 48627 (Fig 6B). A dose-dependent increase in TbpB-specific antibody was observed, with slightly higher reactivity obtained against 41385 TbpB compared to 48627 TbpB. Despite mouse-to-mouse variability, reactivity was observed across the full panel of gonococcal TbpBs (Fig 6C). However, in contrast to the reactivity seen by the immune rabbit serum raised to the bivalent TbpB formulation, minimal reactivity was seen against the meningococcal TbpBs.

**Figure 6:**
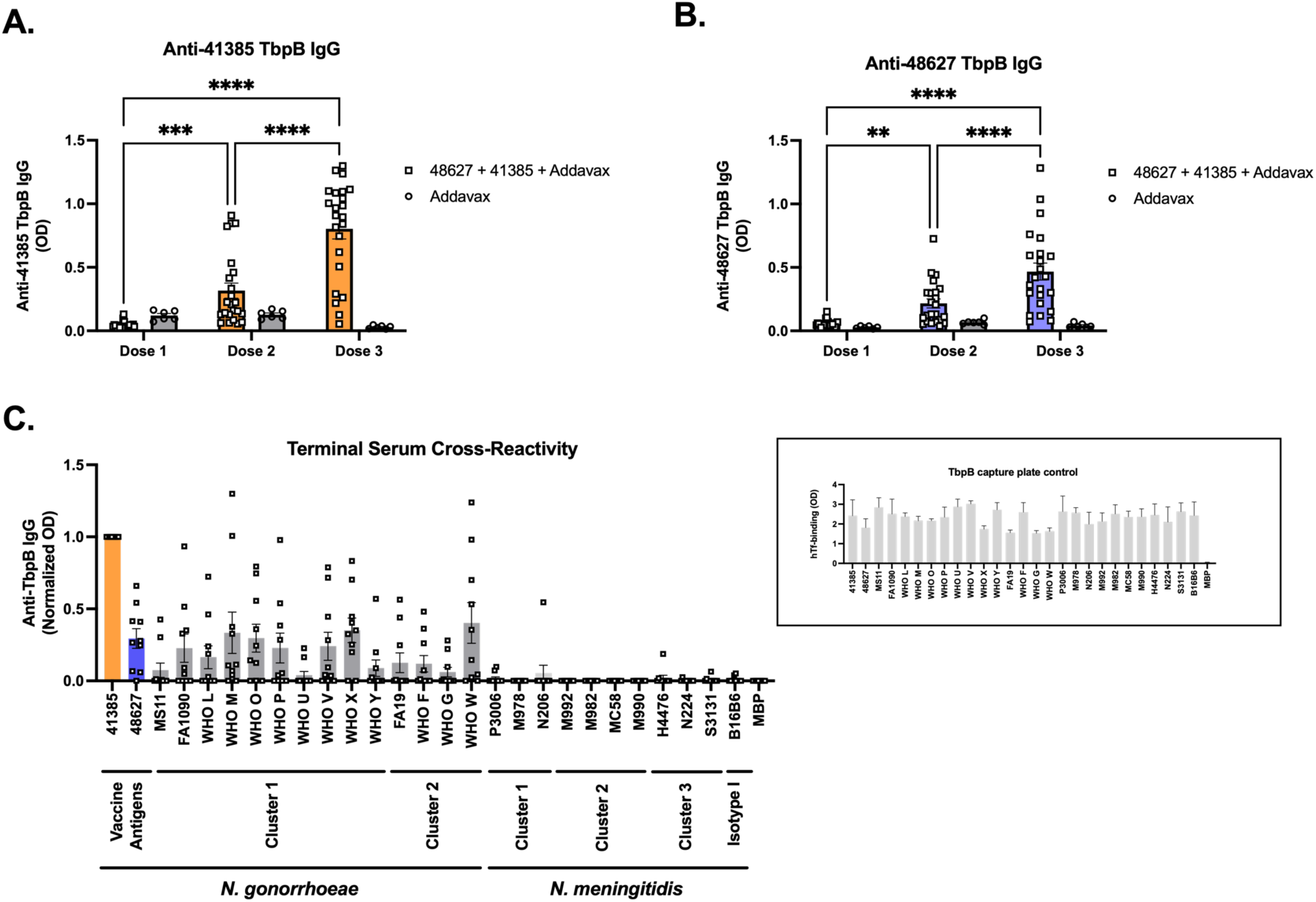
Serum IgG responses in mice to the bivalent TbpB composition. **A, B.** Detection of anti-TbpB IgG in serum of C57BL/6 mice against each immunizing antigen after 1, 2, and 3 doses of the bivalent (Ngo 48627 + Ngo 41385) TbpB vaccine. Separate ELISAs were performed using 41385 TbpB (**A**) and 48627 TbpB (**B**) protein-coated plates to assess the IgG response against each antigen. N=23 for TbpB vaccinated cohort, N=6 for adjuvant controls. Bars represent mean, error bars represent standard error. **C.** Cross-reactivity of terminal serum from bivalent TbpB immunized mice against a broad panel of Neisserial TbpB proteins belonging to different clusters. Optical density (OD) values were normalized with Ngo 41385 TbpB and MBP (empty vector) set at 1 and 0, respectively, with any negative values (signal below empty vector) plotted as 0. N=10 serum samples from mice immunized with bivalent TbpB formulation evaluated. Bars represent mean, error bars represent standard error. Inset, ELISA plate capture control using hTf-HRP binding as a readout to detect folded TbpB. Mean +/- standard deviation of 3 technical replicates. ** represents p<0.01, *** represents p<0.001, *** represents p<0.0001. Non-significant differences in panels A and B not shown. Serum immunogenicity compared between doses by 2 way ANOVA with Sidak’s multiple comparisons test comparing matched samples.

Mice that received the bivalent TbpB formulation were randomized into two groups and infected vaginally with one of two heterologous strains of *N. gonorrhoeae*: either Ngo WHO L, which contains a TbpB 94.0% identical to Ngo 48627, or Ngo WHO F, which contains a TbpB 86.7% identical to Ngo 41385. These strains were selected as being the closest match to the selected immunogens from the WHO panel of strains which was already utilized for murine infections by our group. Mice were followed daily to establish the bacterial burden and duration of colonization. Mice that received the bivalent TbpB composition decreased the median duration of colonization from 12 (Ngo WHO L, Fig 7A, B) or 10 (Ngo WHO F, Fig 7F, G) days for mice that received adjuvant only down to 4 days for both strains, though the difference in clearance was not statistically significant. The recovered CFU from Ngo WHO F was lower in mice that received the bivalent composition compared to mice that received adjuvant only (Fig 7J), however no difference in overall recovered bacteria was seen for Ngo WHO L (Fig 7E).

**Figure 7:**
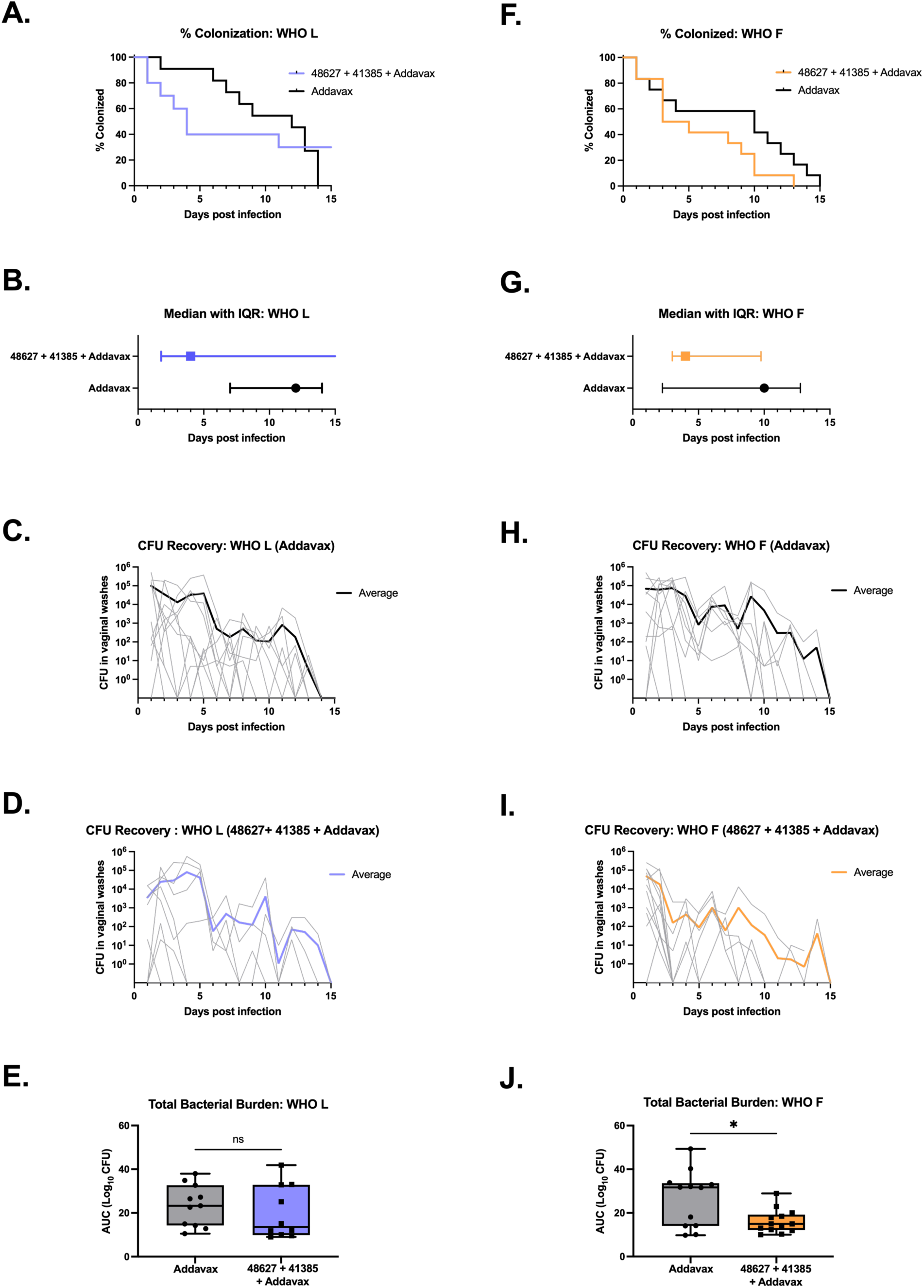
Protection conferred by the bivalent TbpB vaccine formulation against representative heterologous isolates belonging to the two gonococcal TbpB clusters in mice. **A, F.** Percentage of vaccinated and adjuvant controls that remain colonized over a period of 15 days after vaginal infection with a Ngo cluster 1 strain (WHO L) and cluster 2 strain (WHO F), respectively. N=10-12 animals/group. Differences were not significant using Log-rank (Mantel Cox) statistical test. **B, G.** Graph depicting the number of days taken for 50% of animals to clear infection. Error bars depict interquartile range (IQR). **C, D, H, I.** Daily CFU recovery from the lower genital tract of infected mice. Thin grey lines depict individual animals, while thicker lines represent average CFU recovered per day. **E, J.** Box plot showing total bacterial burden in each animal. Bioburden was estimated using area under the curve of Log10 transformed CFU data. *, p<0.05 obtained using two-tailed t-test, ns denotes not significant. Statistical analysis done using GraphPad Prism 10.1.1.

## Discussion

The rational selection of immunogens for inclusion into a vaccine formulation is a fundamental step towards developing a broadly cross-protective vaccine. The use of surface exposed protein antigens as targets for vaccine development allows for the targeting of virulence factors and other proteins essential for survival of the pathogen. However, due to the accessibility of these targets to the immune system, these proteins often evolve to exhibit high levels of antigenic diversity.

In this study, we demonstrate that a bivalent TbpB formulation is sufficient to elicit broad cross-reactivity against a panel of gonococcal TbpBs that encompasses the natural diversity of this protein, and against an equally diverse panel of Ngo strains. Expanding the vaccine formulation to include five gonococcal TbpBs did not further increase the breadth of cross-reaction, indicating that one rationally selected protein variant is sufficient to elicit a response against each of the two Ngo TbpB clusters. This supports the utility of our custom software Navargator to rationally select variants to include in a vaccine formulation. With this, we were able to leverage sequence diversity information by performing clustering using the phylogenetic distance between the variants. This analysis allows us to deterministically define clades, and to select the ‘central’ variant from each.

In rabbits, the bivalent and pentavalent formulations both elicited a response that also recognized a subset of meningococcal TbpBs, however this inter-bacterial species response was not apparent in mice. Regardless, our results suggest that the addition of rationally selected TbpBs originating from Nme strains would likely be more effective to expand this coverage for pan-pathogenic Neisseria protection. The bivalent TbpB formulation also elicited broad bactericidal activity against eight representative gonococcal strains and decreased the median duration of colonization by two heterologous gonococcal strains in the murine lower genital tract colonization model, supporting the protective capacity of this response.

Inclusion of multiple variants of a single immunogen is a strategy that has been successfully utilized in previous vaccine formulations, most notably the bivalent factor H binding protein (fHbp) vaccine (Bivalent rLP2086, also known as ‘Trumenba’, Pfizer), wherein one fHbp variant from each of subfamily A and subfamily B were included as lipidated recombinant proteins^30^. Inclusion of multiple proteins in a vaccine formulation increases the complexity of each phase of development, including process development and process characterization. Therefore, identifying the minimum number of variants in a formulation needed to elicit broad coverage becomes vital in developing a vaccine formulation that is feasible, scalable, and cost effective. As shown here, there was minimal expansion in cross-reactivity, transferrin blocking ability, or bactericidal activity when expanding our two variant protein composition up to five, indicating that there is value in rational selection of immunogens and pre-clinical evaluation to reduce the burden of production.

One limitation to the use of small animal models in pre-clinical vaccine research for host-restricted pathogens is that these animals lack host specific factors that are vital during natural infection. The gonococcus is exquisitely host restricted and able to interact with an array of human factors, including those involved in adhesion (human CEACAMs^31^), immune evasion (human Factor H^32^, human C4bp^33^, human IgA1^34^), and nutrient acquisition (human transferrin^35^, human lactoferrin^36^, human calprotectin^37^, human psoriasin^38^). These factors are not present in small animal models including mice or rabbits, which limit the extent that we are able to accurately model gonococcal infections and the response to vaccination with bacterial proteins able to specifically bind to these human factors. Previous studies using either transgenic mice expressing the human factor of interest (e.g. meningococcal fHbp immunization into mice expressing human Factor H^39^) or studies evaluating homologous receptors in the native host (immunization with *Glaesserella parasuis* TbpB into pigs^24^) have demonstrated that the use of non-binding mutant proteins are superior immunogens compared to wild type versions that maintain their ability to bind to the host protein. This is presumably because the native proteins will complex with their host-derived target when administered as a vaccine component. While beyond the scope of this study, our ongoing efforts are focussed on developing non-binding versions of the TbpBs included in the bivalent formulation that will potentially enhance immunogenicity and safety for use in humans.

The use of rabbits in this study allowed us to gather comparatively large volumes of serum from each animal, however the trade off to that was using only one rabbit per formulation for the initial evaluation due to constraints in space and cost. Animal to animal variability is always a concern as individual differences may skew the interpretation of the resulting data. To strengthen the preliminary results from rabbits, we utilized a follow up experiment in mice with the selected bivalent formulation wherein groups of 24 mice received either the bivalent TbpB formulation or adjuvant (50% v/v Addavax) only, which were then randomized into two challenge groups. Leveraging immunization in two species allowed us to increase the confidence in the results, and lower genital tract infections in mice with two challenge strains has allowed us to interrogate not just the reactivity of immune sera against the proteins or strains of interest but to evaluate potential protection beyond antibody-mediated bacterial clearance. As the immune correlate of protection for gonococcal clearance remains elusive, the use of small animal challenge models remains one of the best tools to understanding vaccine efficacy prior to moving into human clinical trials.

Selection of adjuvants to elicit protection is a vital component of vaccine development. While a broad evaluation of adjuvants was beyond the scope of this study, the use of both aluminum hydroxide gel (Alhydrogel) and Addavax (research grade MF59) in rabbits with the two major immunogens of interest demonstrated that the overall breadth of response was not substantially different between the two formulations. Previous studies with TbpB have shown that formulation with alum-based adjuvants elicited higher levels of anti-TbpB serum IgG versus formulation with Addavax, however strength of the antibody response has not correlated with levels of protection^40^. While Addavax was utilized for the murine protection studies, recent insights by our group^40^ underscore the necessity for additional optimization of the adjuvant to attain robust protection.

Together, the studies shown here demonstrate that broad gonococcal cross-protection can be obtained with a bivalent TbpB formulation selected rationally based on phylogenetic analysis of the diversity of this protein.

## Methods

### Bioinformatic analysis and selection of the TbpBs of interest

To generate the large Neisserial phylogenetic tree, we collected all TbpB sequences from the PubMLST database that were annotated as originating from either *N. meningitidis* or *N. gonorrhoeae*. We removed all sequences shorter than 500 or longer than 750 residues, those containing internal stop codons, and one sequence (ID 21071) that was radically different from all others and is likely a misannotation. We then added several sequences of prototype strains commonly used in various laboratories (MS11, FA1090 and FA19) as well as the WHO reference panel^27^. The signal peptides were removed, and duplicate sequences were filtered out, resulting in 1485 unique total sequences: 1178 from *N. meningitidis* and 307 from *N. gonorrhoeae*. 304 of the 307 sequences from *N. gonorrhoeae* were found in two monophyletic groups, and these were used to build the *N. gonorrhoeae* phylogenetic tree; the 3 outlier sequences were widely spread throughout the large tree and were not included as they were assumed to be misannotated or the result of horizontal exchange from *N. meningitidis*. An additional 21 *N. gonorrhoeae* TbpB sequences were collected from a more recent version of the PubMLST database, and these 325 sequences were used to build the *N. gonorrhoeae* phylogenetic tree.

Sequence alignments were performed using the E-INS-i algorithm with default parameters from MAFFT v7.475 ^41^. The phylogeny was reconstructed with RAxML v8.2.12 ^42^ using the PROTCATLG model of evolution.

To choose representative variants from *N. gonorrhoeae*, we started first with selecting representative variants from two clusters, Ngo 48627 as the large cluster representative, and Ngo 41385 as the small cluster representative. We further used Navargator to identify several potential clustering patterns on the current tree and settled on five clusters as the smallest number that partitioned the tree effectively. This yielded our final list of five representative *N. gonorrhoeae* TbpB sequences: 27464, 48627, WHO Y, 27230, and 41385.

### Cloning, expression, and purification of selected TbpBs

The five representative TbpB variants were codon optimized and synthesized as DNA fragments (GeneArt String, ThermoFisher) with 27 base pairs of complementary sequences matching our custom expression vector (codon optimized sequences available in supplemental Table 1). The inserts were subcloned by overlap extension PCR cloning^43^ downstream and in-frame of an N-terminally hexa-histidine tagged maltose binding protein (MBP) and a cleavable TEV linker sequence. *Escherichia coli* MM294 was used to amplify the vectors and sequence verification was done by in-house Sanger sequencing (The Center for Applied Genomics, The Hospital for Sick Children). Protein expression was carried out in *E. coli* Overexpression^TM^ C43 DE3 cells (Sigma Aldrich), freshly heat-shocked transformants were recovered in Luria-Bertani (LB) broth for 1 hour, shaking at 175 RPM and 37°C, before selection by additional LB broth (3 mL) supplemented with ampicillin (100 μg/mL) and grown for a further 3 hours. The starter culture was inoculated into 6 L of ZY autoinduction media supplemented with NPS and 5052 and ampicillin (50 μg/mL)^44^ at a 1:1,000 dilution with the culture flasks shaken at 175 RPM at 37°C for 18 hours and then at 20°C for additional 12 to 16 hours.

Cells were harvested by centrifugation at 5,000 x g for 30 minutes at 4°C, resuspended in binding buffer (50 mM Tris pH 8.0, 300 mM NaCl) supplemented with DNase, lysozyme, and protease inhibitors (cOmplete^TM^ Mini, Roche) and either lysed by sonication (S-450 Sonifier, Branson) or homogenization (EmulsiFlex-C3, Avestin). The resulting lysate was clarified by centrifugation at 16,000 x g for 90 minutes at 4°C, followed by 0.22 μm syringe filtration (Millipore). Clarified lysate was then recirculated and captured by nickel Sepharose affinity chromatography (HisTrap excel, Cytiva). The affinity column was then mounted onto an ÄKTA Purifier 100 (GE Healthcare), washed, and eluted with imidazole containing buffer (50 mM Tris pH 8.0, 300 mM NaCl, 300 mM imidazole pH 7.4). Eluted fraction was dialyzed overnight into exchange buffer (50 mM Tris pH 8.0, 150 mM NaCl) and sample purity assessed by 12% SDS-PAGE analysis and concentration by NanoDrop absorbance at 280 nm (ThermoFisher); TEV protease, purified in-house, was added to separate the MBP fusion partner from TbpB. The samples were then exchanged into a low salt buffer (50 mM Tris pH 8.0, 10 mM NaCl) and loaded onto an anion exchange column (HiTrapQ FF, Cytiva) and the fraction containing TbpB concentrated and passed through a size exclusion column (HiPrep 26/60 Sephacryl S-200 HR, Cytiva) to further remove any contaminants and exchange into the final sample buffer (50 mM HEPES, 50 mM NaCl). A final polishing step with a strong anion exchange column (MonoQ 5/50 GL, Cytiva) was used to remove lipopolysaccharide and the final sample was concentrated to 5 mg/mL in aliquots, flash frozen by liquid nitrogen, and store at -80°C until use.

### Rabbit immunizations

Rabbits were provided food and water *ad libitum* and handled humanely under animal use protocol 20012577, approved by the Animal Care Committee at the University of Toronto. New Zealand female rabbits (Charles River) were obtained at approximately three months of age and acclimated to the facility for one week. Rabbits 1-5, 8, and 9 (Table 2) received an initial dose of 100 μg of protein formulated with the respective adjuvant (38.5 μl of 2% Alhydrogel per 500 μl dose, Sigma Aldrich, cat. A8222 or 50% (v/v) Addavax, InvivoGen, cat. vax-adx-10) in 500 μg total volume, immunized subcutaneously. Two booster doses were given containing half the amount of protein as the initial vaccine at 3 and 6 weeks post first dose. Rabbit 6 received the bivalent TbpB composition which was composed of 100 μg of each TbpB (dose 1) or 50 μg of each TbpB (doses 2 and 3) for formulations containing 200 μg and 100 μg total protein respectively, while Rabbit 7 received the pentavalent formulation composed of 50 μg of each TbpB (dose 1) or 25 μg of each TbpB (doses 2 and 3) for formulations containing 250 μg and 125 μg total protein, respectively (Table 2). Bleeds were collected 2.5 weeks after each dose and at euthanasia (two weeks after the third dose).

### Protein ELISAs

Protein ELISAs were performed as previously described^23^. Briefly, TbpBs of interest were cloned into a custom His-Bio-MBP-Tev vector and transformed into *E. coli* strain ER2566. *E. coli* strains were grown overnight in 10 mL cultures of ZYP-5052 autoinduction media containing 100 𝜇g/mL ampicillin and grown overnight at 37°C with shaking. Lysates were obtained by centrifuging the *E. coli* cultures (3,000 × g for 5 minutes) and resuspending the pellets in buffer (50mM NaHP_2_PO_4_, 300mM NaCl, 10mM imidazole, pH 8.0). 1 mm glass beads were added to the suspension and cells were disrupted by shaking in a cell disruptor. Homogenate was centrifuged for 20 min at 4°C at 14,000 x g. Supernatant was diluted 1:5 in PBS and added to 384 well ELISA plates (VWR, cat. CA62409-064) that had been previously coated with 1 𝜇g/mL NeutrAvidin (Thermo Fisher, cat. A2666, coated overnight at 4°C) and blocked with 5% bovine serum albumin (BSA, BioShop Canada, cat. ALB001). Protein lysate was coated for 2 hours at room temperature, washed, and then plates were blocked again with 5% BSA for 1 hour. Serum was added overnight at 4°C at a 1:8,000 dilution for rabbit sera and at a 1:10,000 dilution for mouse sera. The following day, plates were washed, the relevant secondary antibody was added (1:10,000 dilution of goat anti-mouse IgG H&L (HRP) (Abcam, cat. ab6789) or 1:10,000 dilution of Peroxidase AffiniPure goat anti-rabbit IgG (H&L) (Jackson ImmunoResearch, cat. 111-035-144)) and incubated at room temperature for two hours. Plates were washed and developed with KPL SureBlue TMB Microwell Peroxidase Substrate (SeraCare, cat. 5120-0077). Reactions were quenched with 2 N H_2_SO_4_ and read at 450/570nm.

### Gonococcal strains

**A.** *N. gonorrhoeae* strains used for these studies include laboratory strains FA19, MS11, and FA1090, the WHO reference strains (obtained from Public Health England culture collection)^27^, clinical isolates obtained from Public Health Ontario which consists of both mucosal and disseminated strains (described in supplemental Table 2), and a collection of gonococcal strains acquired from endocervical swabs from female sex workers in Nairobi, Kenya (Kenyan Ngo Collection, described in supplmental Table 2)^28,45^.

### Whole cell ELISAs

Whole cell ELISAs were performed with a panel of representative strains of *N. meningitidis* and *N. gonorrhoeae.* Strains were grown overnight on GC agar supplemented with Kellogg’s and 10 𝜇M deferoxamine mesylate salt (Sigma, cat. D9533) at 37°C with 5% CO_2_ and then restreaked onto several fresh GC agar supplemented with Kellogg’s and 10 𝜇M deferoxamine mesylate salt for another overnight growth. Bacteria were resuspended in PBS and heat killed at 55°C for a minimum of 60 minutes. 384 well ELISA plates were coated with killed bacteria in PBS at an OD_600_ of 0.2 and dried in a biosafety cabinet until heat killing of bacteria was confirmed. Plates were blocked with 5% BSA and ELISAs were performed as described above.

### Transferrin blocking

Transferrin blocking assays were done using the protein-based ELISA protocol described above with rabbit serum added in a 1:10 dilution; and instead of probing with goat-anti-rabbit IgG, horse radish peroxidase conjugated hTf (HRP-hTf, Rockland, cat. 009-0334) was added at a dilution of 1:1,000 and incubated at room temperature for 30 minutes. Plates were developed as above, and a decrease in signal indicated blocking of transferrin binding. Signal was compared to transferrin binding to the protein in question with no serum present.

### Serum bactericidal assays

Serum bactericidal assays (SBA) were performed with 8 strains of *N. gonorrhoeae* with varying concentrations of pooled normal human serum (NHS) (Sigma, cat. H4522, Lot #SLCD1946), each using the highest concentration of NHS (range 2.5% to 15%) that the strain was intrinsically resistant to in the absence of any antibody source. Strains were grown overnight on GC agar supplemented with Kellogg’s at 37°C with 5% CO_2._ The next day strains were restreaked onto GC agar supplemented with Kellogg’s and 100 𝜇M deferoxamine mesylate salt (Sigma, cat. D9533) for 4 hours. Bacteria were collected off the plates using Dacron swab, resuspended in PBS++, and OD_550_ was measured. Assay was set up in 40 𝜇l total volume using ∼1000-2000 bacteria suspended in RPMI, NHS, and 2-fold serial dilutions of heat inactivated rabbit serum and incubated at 37°C with 5% CO_2_ for 60 minutes and then plated. The highest dilution at which 50% killing was observed relative to no antibody control (NHS only) at 60 minutes was reported as the SBA titre.

### Mouse immunizations and genital tract challenge

Mouse studies were performed under the animal use protocol 20011775, approved by the Animal Care Committee at the University of Toronto. Female C57BL/6 mice (4-5 weeks of age at time of purchase, Charles River) were acclimated to the animal facility for one week where they received water and rodent chow *ad libitum* and were housed in specific pathogen free conditions. Mice were randomly assigned to receive either the bivalent TbpB formulation (12.5 𝜇g 48627 TbpB, 12.5 𝜇g 41385 TbpB in sterile PBS, formulated with 50% v/v Addavax (InvivoGen) in 100 𝜇l total volume) or adjuvant alone (50% v/v Addavax in PBS). Mice were immunized three times at three-week intervals via intraperitoneal injection. Vaccines were well tolerated, and no adverse reactions were noted. Saphenous bleeds were performed two to three weeks after each dose and serum was stored at -20°C until analysis.

Two weeks after the final dose, mice were challenged via the lower genital tract challenge model^46^. Briefly, the stage of the estrus cycle was tracked via evaluation of cell morphology of vaginal lavage samples under light microscopy. When mice entered diestrus (denoted as day -2 relative to infection), they were given β-estradiol (0.5 mg in 200 𝜇l sub-cutaneous, Sigma-Aldrich, cat. E4389) to lock them in estrus and antibiotics (intraperitoneal 0.6 mg vancomycin (BioShop Canada, cat. VAN990.5) and 2.4 mg streptomycin (BioShop Canada, cat. STP101.5), as well as 0.04 g/100mL trimethoprim (Sigma-Aldrich, cat. T7883) in the drinking water *ad libitum*) to control overgrowth of commensal flora. Mice received two doses of injectable antibiotics the following day (day -1 relative to infection). On day 0, mice were infected intra-vaginally with approximately 10^7^ CFU of either *N. gonorrhoeae* WHO L or *N. gonorrhoeae* WHO F (obtained from Public Health England^27^, streptomycin resistance induced). Mice received two additional doses of β-estradiol (one on the day of infection and another on day +2 relative to infection) and daily antibiotic injections for the duration of the study. Vaginal lavage samples (15 𝜇l PBS++ diluted with 60 𝜇l) were collected daily and dilutions were plated on GC agar supplemented with Kellogg’s and VCNT (vancomycin, colistin, nystatin, and trimethoprim) inhibitor (BD BBL^TM^ VCNT Inhibitor (Thayer and Martin), Fisher Scientific, cat. B12408) for selection of *N. gonorrhoeae.* Mice were considered no longer colonized when a minimum of three consecutive lavages had no recovery of *N. gonorrhoeae.* Group sizes for each vaccine group and challenge strain can be found in Table 3.

**Table 3:** Group sizes for the murine lower genital tract colonization study broken down by challenge strain.

| Immunization Group | Number immunized (n) | Challenge strain | Number infected (n) |
| --- | --- | --- | --- |
| Bivalent TbpB | 24 | WHO L | 10 |
|  |  | WHO F | 12 |
| Addavax Only | 24 | WHO L | 11 |
|  |  | WHO F | 12 |

Upon completion of the infection study, mice were humanely euthanized by CO_2_ overdose. Terminal serum was collected by cardiac puncture and stored at -20°C until analysis.

### Graphing and statistical analyses

All graphing and statistical analyses were performed in Prism 10.1.1. Phylogenetic tree figures were created using Navargator^22^.

### Author contributions

JEF conceptualized the rabbit immunization study; conceptualized, performed, analyzed and interpreted the murine infection study and related serological analysis; and wrote the original manuscript. EAI conceptualized, performed, analyzed, and interpreted rabbit and mouse cross-reactivity ELISAs and serum bactericidal assays; performed and interpreted the murine infection study and related serology; contributed to writing and editing the manuscript. DMC performed the bioinformatics analysis and contributed to writing and editing the manuscript. DN designed constructs and performed the protein purification for the rabbit immunizations; contributed to writing and editing the manuscript. NA performed the protein purification for the murine immunizations, provided technical support for the murine challenge study, and edited the manuscript. EGC performed the murine infection study and related serology, and edited the manuscript; JZ and JL performed the murine infection study, and edited the manuscript. ABS, TFM, and SDG provided conceptualization support, supervision, and edited the manuscript.

## Acknowledgements

The authors would like to acknowledge Dr. Vanessa Allen (Public Health Ontario) for generously sharing recent gonococcal isolates collected in Ontario, Canada, from between 2015 and 2018, Dr. Gursonika Binepal for helping prioritize which Public Health Ontario strains to include in the present study, Ethan Gray-Owen for technical support during cloning and ELISA optimization, and Dr. Nelly Leung for general laboratory support. The authors would also like to thank the animal support staff at the Division of Comparative Medicine at the University of Toronto for technical and welfare support for both the rabbit and mouse studies presented here, including performing rabbit immunizations and blood sampling. The authors appreciate the financial support received for the research, authorship, and/or publication of this article. This research was supported by National Institute of Health funding #R01-AI125421-01A1 and #1RO1AI141229. SDG and TFM are supported by the Canada Research Chair Program.

## Notes

### Competing Interest Statement

JEF, EAI, DMC, DN, NA, ABS, TFM and SDG control intellectual property related to transferrin receptor-based vaccines for use in humans and animals. No payments have been received related to the work described herein.

