## Supplemental Tables for "Rational selection of TbpB variants elucidates a bivalent vaccine formulation with broad spectrum coverage against *Neisseria gonorrhoeae*"

Table S1: Codon optimized sequences of the five representative TbpB variants

**>Ngo48627**

CAACGAAAACCTGTATTTTCAGGGATCCCCTGCACCGAAATATCAGGATGTTCCGAGCAAAAAACCGGAAGCACGTAAAGATCAAGGTGGTTATGGTTTTGCCATGCGTTTTAAACGTCGTAATTGGTATCCGCCTAGCAATCCGAAAGAAAATGAAATTCGTCTGAGCGAAGGTGATTGGGAACAGACCGATAATGGTGATATCAAAAAACCCAGCAAGCAGAAGGATATCATTAACGCACTGAGCGGTAATGGTGGTGAACTGCTGCAGGATAGCAGCCAGCGTGGTAAAGGTATTAGCAAAGTTACCGATCACCACGACTTTAAATACGTTTGGAGCGGCTTTTTCTATAAGCAGATTGGTAACACCGTGAAAAAAAGCGGTAGCAGCATTACCGAAGCGCGTAATGGTCCGGATGGCTATATCTTCTATAAAGGTAAAGATCCGAGCCGTGAACTGCCGGTTCTGGGTAGCGTTGAATATAAAGGCACCTGGGATTTTCTGACCGATGTTCGTGTTAATCAGAAATTTACCGATCTGGGTAGTGCAAGCACCAAAAGCGGTGATCGTTATAGCGCATTTAGCGGTGAACTGGATTATATCGTGAAAAAGCAAGAGGATAAAAAAGAAAAACACAAAGGTCTGGGTCTGACCACCGAAATTACCGTTGATTTTGAGAAAAAAAACCTGAACGGCAAACTGATCAAGAACAACTATGTCATTAACAACAACAACGGCGACGACCAGAACAAATATACCACCGAGTATTATACCCTGGATGCAACCCTGCGTGGTAATCGTTTTAGCGGTAAAGCAACCGCAACCGATAAAAGCAGTGATGGTCAGGCAAAACAGCATCCGTTTGTTAGCGATAGCAGCAGCCTGAGCGGTGGTTTTTTTGGTCCGCAGGGTGAAGAACTGGGTTTTCGCTTTCTGAGTGATGATGGTAAAGTTGCAGTTGTTGGTAGCGCAAAAACCAAAGATAAAAATGCCAATGGTAATACCGCAGCAGCAGGTACAGCCGGTGCAGCCGGTATGAGCAGCGAAGATACCCGTCTGACAACCGTTCTGGATGCAGTTGAACTGACACTGGATGGCAAAAAAATCAAAGATCTGGATAACTTTAGTGACGCAACCCGTCTGGTTGTTGATGGTATTATGATTCCGCTGCTGCCGAATGATAGCGGTAGCGGTGGTAGCCAGGCAGATAAAGGCAAAAATGGTGGCACCGCATTTACCTATGAAACCACCTATACACCGGAAAGCTATACCCCTGAAAGCGACAAAAAAGATACCAAAGCAGGCACCGCAGCAAATGGTGTTCAGACCGTTAGCAATACTGCCGGTGGCACCAGCGGTAAAACCAAAACACATTATAAAGTTCAGGTGTGTTGCAGCAATCTGAACTATCTGAAATATGGTCTGCTGACCCGTGAAAATAACAATAGCGTTATGCAGGCAGTGAAAAATAGCAATCGTACCGCAGATCGCACCGCACAGGGTGCACAGAGCATGTTTCTGCAGGGCGAACGTACCGATGAAAAAGAAATTCCGAAAGATGAGAACGTGGTGTATTTAGGTACATGGTATGGTCATATTGCAACCAATGGTACAAGCTGGACACGTGAAGCAAGCAATCAAGAAAATGGTAATCGCGCAAAATTCGATGTGAACTTCAAAGACAAACGCATTACCGGTACACTGACCGCAGAAAATCGTAGCGAAGCAACCTTTACAATTGATGCCATGATTGATGGCAATGGCTTTAAAGGTACAGCAAAAACCGGCAATGATGGCTTTGCACCGGATCAGAATAGCAGCACCGGCACCTATAAAGTGCATATTGCCAATGCCGAAGTTCAAGGTGGCTTTTATGGTCCTAATGCGGAAGAATTAGGTGGTTGGTTTGCATATCCTGGTAATGGCCAGGCCAAAAATGCACAGGCAAGCAGCGGTAGTGGTAATAGTGCAGGTAGCGCCACCGTTGTTTTTGGTGCAAAACGTCAGCGTCTGGTGAAATAAAAGCTTCTGCCTGGCGGCCTTTTGCGCC

**>Ngo27230**

CAACGAAAACCTGTATTTTCAGGGATCCGCAGCACCGAAATATCAGGATGTTCCGAGCAAAAAACCGGAAGCACGTAAAGATCAAGGTGGTTATGGTTTTGCCATGCGTTTTAAACGTCGTAATTGGCATCGTATGGCCAATGAAAATGAAGTGAAACTGAATGAAAGCGATTGGGAACAGACCGATAATGGCAACATTAAAGAACCGAGCAAACAGAAAAGCATTATTGATGCACTGACCGGTAATGATGGTGAAACCCTGCAGGATAGCAGCCAGCAAGAAGGTATTAGCAAAGTTACCGGTTATCACGACTTCAAATATGTGTGGTCAGGCTTCTTCTATAAACACATTCGTACCAAAAGCGAAACCATTGATGGTAAAGTGACCGTTCGTAGCGGTCCGGATGGCTATATTTTCTATAAAGGCACCGATCCGAGCCGTAAACTGCCGGTTAGCGGTAAAGTTATGTATAAAGGTACATGGGATTTTCTGACCGATGTGAAAGCCAATCAGAAATTTACCGATCTGGGTAATGCAAGCGCAAAACCGGGTGATCGTTATAGCGCATTTAGCGGTGAACTGGATTACATCGTGAAAAAAGAGGACGATAAAAAAGACGGTCATGTTGGTCTGGGTCTGACCACCGAAATTACCGTTGATTTTGGTAAAAAAACCCTGAGCGGCAAACTGATCAAAAACAACATGGTGATTAATAACGGTGATGAACCGACCACGCAGTATTATTCACTGGAAGCACAGGTTACCGGCAATCGTTTTAATGGTAAAGCAATTGCAACCGACAAACCGAAAGCAAACGAAACCAAAGAACATCCGTTTGTTAGCGATAGCAGCAGCCTGAGCGGTGGTTTTTTTGGTCCGAAAGGTGAAGAACTGGGTTTTCGCTTTCTGAGTGATGATGGCAAAGTTGCAGTTGTTGGTAGCGCAAAAACAAAAGATGAAACCGCAAGCAGCGGTGGCACCAGCGGTGGTGCAAGCGTTAGCGCAAGCGGTGGTACAACCGGTACACCGAGCGAAAATAAACTGACCACAGTTCTGGATGCAGTTGAACTGACACCGGATGGCAAAAAAATCAAAGATCTGGATAACTTTAGCAACGCAGCACAGCTGGTTGTTGATGGTATTATGATTCCGCTGCTGCCGACCGAAAGCGGTAATGGTCAGGCAGATAAAGGTGAAAATGGTAAAACCGCCTTCATCTATGAAACCACCTATACACCGGAAAGCGACAAAAAAGATACCCAGACCGGCATGGCAACCAATGGTGTTCAGACCGTTAGCAATACCGCAGGCGGTACAAGCGGTAAAACCAAAACACATTATAAAGTTCAGGCCTGTTGCAGCAATCTGAACTATCTGAAATATGGTCTGCTGACCCGTAAAAATAGCGAAAGCGCAATGCAGGCAGGCGAAAGCAGCAGTCGTACCGCAGTGCAGACCGCACAGGGTGCACAGAGCATGTTTCTGCAGGGTGAACGTACCGATGAAAAAGAAATTCCGAAAGATGGCAACGTGGTTTATTTAGGCACCTGGTATGGTCATATTGCAATTAATGGCACCAGTTGGACCCGTGAAGCAAGCAATCAAGAAAATGGCAATCGTGCCAAATTCGACGTGAACTTCAAAGACAAAAAGATTACCGGTACGCTGACCGCAGCAAATCGTCAAGAAGCAACCTTTACAATTGATGCCATGATTGAAGGCAATGGCTTTAAAGGTACGGCAAAAACCGGTGATGGTGGCTTTGCACCGGATCAGAATAATAGCACCGGCACACATAAAGTGCATATTGCCGAAGCAAAAGTGCAAGGTGGCTTTTATGGTCCTAATGCCGAAGAATTAGGTGGTTGGTTTGCATATCCTGGCAATGGCCAGGCCGAAAATGCACAGACCAGCAGTGGTAATGGTAATAGCGCAGGTAGCGCCACCGTTGTGTTTGGTGCGAAACGCCAAGAACTGGTGAAATAAAAGCTTCTGCCTGGCGGCCTTTTGCGCC

**>Ngo27464**

CAACGAAAACCTGTATTTTCAGGGATCCGCAGCACCGAAATATCAGGATGTTCCGAGCAAAAAACCGGAAGCACGTAAAGATCAAGGTGGTTATGGTTTTGCCATGCGTTTTAAACGTCGTAATTGGTATCGTGCCACCAATGAAAATGAGGTGAAACTGAAAGAAAGCGATTGGGAACAGACCGATGATGGCGAAATCAAAAATCCGTTCAAGCAGAAGAACATCATTAACGCACTGCCTGGTAATGAAGGTGAACTGCTGCAGGATAGCAGCCAGCAAGGTAAAGGTATTAGCAAAGTTGGTGATCATCACGATTTCAAGTATGTTTGGAGCGGCTTTTTCTATAAACGCATTGAAATCACCACCAAAAAAAACGAAAGCAACAAGATTATTGAAGCCCGTAGCGGTCCGGATGGCTATATCTTCTATAAAGGTGGTAATCCGAGCCGTAAACTGCCGGTTAGCGGTGAAGTTACCTATAAAGGCACCTGGGATTTTCTGACCGATGTTAAAGCAAATCAGCGTTTTACCGATCTGGGTAATACCAGCACCAAAAGCGGTGATCAGTATAGCGCATTTAGTGGTGAACTGGATTACATCGTGAAAAAAGAAGAGGATAAAAAAGAAAAACATAAAGGCCTGGGTCTGACCACCGAAATTACCGTTGATTTTGGTAAAAAAACCCTGAACGGCAAGCTGATCAAAAACAACAAACTGATTAACAACAACGATGAACCGACCACGCAGTATTATACCTTTGATGCAACCCTGCGTGGTAATCGTTTTAGCGGTAAAGCAACCGCAACCGACAAAAAAGAGAATGAAACCGGTCAGCATCCGTTTGTTAGCGATAGCAGCAGCCTGAGCGGTGGCTTTTTTGGTCCGCAGGGTGAAGAACTGGGTTTTCGCTTTCTGAGTGATGATGGTAAAGTTGCAGTTGTTGGTAGCGCAAAAACCAAAGATAATACCGCAAATGGCAATCCGGCAGTGTCAAGCGGTGCCGGTGCAGCAGCAATGCCGAGCGAAACAGGTCTGACAACCGTTCTGGATGCAGTTGAACTGACCCTGGATGGCAAAGAAATTAAGAACCTGGATAACTTTTCAGATGCGACCCGTCTGGTTGTTGATGGTATTATGATTCCGCTGCTGAGCACCGAATCAGGTGATGGTCAGGCAGATAAAGGTAAAAATGGTGGCACCGATTTTACATATACCACCACCTATACCACAACGTATACACCGGAAAGCGATAAAAAGGATACCAAAGCACAGACCGGTGCAGTTGGTATGCAGACCGCACCGGGTGCAGCGGGTGTTAATGGTGGCCAGGCAGGCACCAAAACCTATGAAGTTGAAGCATGTTGTAGCAACCTGAACTATCTGAAATATGGTATGCTGACCCGCAAAAATAGCGAAAGCGCAATGCAGGCTGGTAAAAATAGCTCACAGGCAGATGCCAAAACCAAGCAGATTGAACAGAGCATGTTTCTGCAGGGCGAACGTACCGATGAAAAAGAAATTCCGAAAGATGAGAACGTGGTGTATTTAGGTACATGGTATGGTCATATTGCAACCAATGGCACCAGCTGGACCCGTGAAGCAAGCAATCAAGAAAATGGTAATCGCGCAAAATTCGATGTGAACTTCAAAGACAAACGTATTACCGGTACACTGACCGCAGAAAATCGTAGCGAAGCCACCTTTACCATTGATGCCATGATTGATGGTAATGGCTTTAAAGGTATGGCCAAAACCGGTAATGGTGGTTTTGCACCGGATCAGAATAGCAGCACCGGTACGCATAAAGTTCATATTACCAGCGCAGCAGTGCAAGGCGGTTTTTATGGTCCGAAAGCCGAAGAATTAGGTGGTTGGTTTGCATATCCTGGCAATGGCCAGACCAAAAATGCACAGGCAAGCAGTGGTAATGGTAATAGTGCAGGTAGCGCCACCGTTGTGTTTGGTGCCAAACGTCAAGAACTGGTGAAATAAAAGCTTCTGCCTGGCGGCCTTTTGCGCC

**>NgoWHO_Y**

CAACGAAAACCTGTATTTTCAGGGATCCCCTGCACCGAAATACAAAGATGTTCCGAGCAAAAAACCGGAAGCACGTAAAGATCAAGGTGGTTATGGTTTTGCCATGCGTTTTAAACGTCGTAATTGGTATCCGCCTAGCAATCCGAAAGAAAATGAAATTCGTCTGAGCGAAGGTAATTGGGAACAGACCGATGATGGTGAAATCAAAACCCCGAGCAAACAGAAAAACATTATTAACGCACTGAGCGGTAATGAAGGTGTTAGCCTGCAGGATAGCAGCCAGCAAGGTGAAGGTATTAGCAAAGTTACCGATCACCACGACTTTAAATACGTTTGGAGCGGCTTTTTCTATAAACGCATTGGTATCACCACCAAAAAAGATGATCTGAGCAACAAGATTATCGAAGCCCGTAATGGTCCGGATGGCTATATCTTCTATAAAGGCACCGATCCGAGCCGTAAACTGCCGGTTAGCGGTAGCGTTGAATATAAAGGTACATGGGATTTTCTGACCGATGTGAAAGCCAATCAGAAATTTACCGGTCTGGGTAATACCAGCACCAAAAGCGGTGATCGTTATAGCGCATTTAGCGGTGAACTGGATTACATCGTTAAAAAAGAAAGCGACAAAAAGGATGGTCATGTTGGTCTGGGTCTGACCACCGAAATTACCGTTGATTTTGGTAAAAAAACCCTGAGCGGCAAACTGATCAAAAACAACATGGTGATTAATAACGGTGATGAACCGACCACACAGTATTATTCACTGGAAGCACAGGTGACCGGTAATCGTTTTAATGGTAAAGCAATTGCAACCGACAAACCGAAAGTGAACGAAACCAAAGAACATCCGTTTGTTAGCGATAGCAGCAGCCTGAGCGGTGGTTTTTTTGGTCCGCAGGGTGAAGAACTGGGTTTTCGCTTTCTGAGCCATGATAATAAAGTTGCAGTTGTTGGTAGCGCCAAAACCAAGGATAAAAATGCAAATGGTAATACCGCAGCAGCAGGTACAGCCGGTGCAGCCGGTATGAGCAGCGAAGATACCAAACTGACCACAGTTCTGGATGCAGTTGAACTGACACCGGATGGCAAAAAAGTTAAAAACCTGGATAACTTTAGTGACGCAACCCAGCTGGTTGTTGATGGTATTATGATTCCGCTGCTGCCGACCGAATCAGGTAATGGTCAGGCAGATAAAGGTGAAAATGGTAAAACGGCCTTCATCTATGAAACCACCTATACACCGGAAAGCGATAAGAAAGATACCCAGACAGGCATGGCAACCAATGGTGTTCAGACCGTTAGCAATACTGCCGGTGGCACCAGCGGTAAAACCAAAACACATTATAAAGTTCAGGCCTGTTGCAGCAATCTGAACTATCTGAAATATGGTCTGCTGACCCGTGAAAATAGCAATAGCGTTATGCAGACCGTTCGTAATAGCAGTCAGGCAGCAGCCCGTACCGAACAGGGTGCACAGAGCATGTTTCTGCAGGGCGAACGTACCGATGAAAAAGAAATCCCGAAAGAACAGAAAGTGGTGTATTTAGGCACCTGGTATGGTCATATTGCAGCCAATGGTACAAGCTGGACCGGCAAAGCAAGCGATCAGCAGAGTGGTAATCGTGCAAAATTTGACGTGAACTTCAAGGACAAAAAAATCACAGGCACCCTGACCGCAGCAAATCGTCAGGCCGAAACCTTTACAATTAGCGGTATGATTGATGGCAATGGTTTTGAAGGCACCGCAAAAACCGGTAATGGTGGTTTTGCACTGGATGCAAATAATACAGCAGCAACCCATAAAGCACATATTGCCGAAGCAAAAGTTCGTGGTGGCTTTTATGGTCCTAATGCCGAAGAATTAGGTGGTTGGTTTGCATATCCTGGCAATGGCCAGGCAAAAAATGCCCAGGCAAGCAGTGGTAATGAAAATAGTGCAGGTAGCGCAACCGTTGTGTTTGGTGCAAAACGTCAGCAGCTGGTTCAGTAAAAGCTTCTGCCTGGCGGCCTTTTGCGCC

**>Ngo41385**

CAACGAAAACCTGTATTTTCAGGGATCCCCTGCACCGAAATATCATGATGTTCCGAGCAAAAAACCGGAAGCACGTAAAGATCAAGGTGGTTATGGTTTTGCCATGCGTTTTAAACGTCGTAATCGTCATCCGATGGCAATGCCGAAAGAAAATGAAGTGAAACTGAAAGATGATGACTGGGAAGCAACCGGTCTGCCTGGTGATCCGAAAGATCTGCCAGGTCGTCAGAAAAGCGTTATTGATGAAGTTAGCGCCAATGGCAACAACGATATCTATTTTAGCCCGTATCTGAAACCGAGCAATCATCAGAATAGCAGCATTAATGGTAGCGCAAATCAGCCTCGTAACGAGGTTAAAGATTACAAGAACTTCGAGTATGTGTATAGCGGCTGGTTCTATAAACATGCCAAACCGATTATTGATGGCACCCAGAATAAACTGCAGCAGGGTGATGATGGCTATATCTTTTATCATGGTAAAGATCCGAGCCGTCAGCTGCCTGCAAGCGAAAAAGTTATCTATAAAGGTGTGTGGCACTTTGTGACCGATACCAAACGTGGTCAGAAATTTAACGATATTCTGGAAACCTCCAAAAAACAGGGTGATAGCTATAGCGGTTTTAGCGGTGATGAAGGTGAAACCACCAGTAATCGTACCGATAGTAATCTGAACGATAAACATGAAGGCTATGGCTTTACCAGCGATCTGGAAGTTGATTTCAACAACAAAAAACTGACCGGCAAACTGATTCGCAATAACAAAGTTACCAATGCAGCAGCCAGTGATGGTTATACCACCGAGTATTATACCCTGGATGCAACCCTGCGTGGTAATCGTTTTAGTGGTGAAGCCACCGCAACCGATAAACCGGAAAATGGTAAAAGCAAACAGCATCCGTTTGTTAGCGATAGCAGCAGCCTGAGCGGTGGTTTTTTTGGTCCGCAGGGTGAAGAACTGGGTTTTCGCTTTCTGAGTGATGATAATAAAGTTGCCGTTGTTGGTAGCGCCAAAACCAAAGATAAAAATGCCAATGGTAATACCGCAGCAGCAGGTACAGCCGGTGCAGCCGGTATGAGCAGCGAAGATACGAAACTGACCACCGTTCTGGATGCAGTTGAACTGACACCGGATGGTAAAAAAGTTAAAAACCTGGATAACTTTAGCGACGCAACCCGTCTGGTTGTTGATGGTATTATGATTCCGCTGCTGCCGACCGAAAGCGGTAATGGTCAGGCAGATAAAGGTAAAAACGGTAAAACCGCCTTCATCTATGAAACCACATATACACCGGAAAGCGACGAAAAAGATACCCAGACCGGCATGGCAGCAAATGGTGTTCAGACCGTGAGCAACGCAGCCGGTGGCACCAGCGGTAAAACAAAAACCCATTATGAAGTTCAGGCCTGTTGCAGCAATCTGAACTATCTGAAATATGGTCTGCTGACCCGCAAAACCGCAGATAATACAATGGGTAGCGGTAGCGGTTCACAGGCAGCAGCACAGACCGCACAGGGTGCACAGAATATGTTTCTGCAGGGCGAACGTACCGATGAAAAAGAAATTCCGAAAGAACAGAACGTGGTGTATTTAGGCACCTGGTATGGTCATATTGCCGCAAATGGTACAAGCTGGACCGGTAATGCAAGCGATCAGCAGAGCGGTAATCGTGCACGTTTTGGTGTTAACTTTAAAGACAAAACCATTACAGGCACCCTGACCGCAGAAAATCGTAGCGAAGCAACCTTTACAATTAACGCCATGATTGATGGTAACGGCTTTAAAGGCACCGCAAAAACAGGTAATGATGGCTTTGCACCGGATCAGAATAATAGCACCGTTACACATAAAGTGCATATTGCCAATGCGGAAGTTCAAGGTGGCTTTTATGGTCCTAAAGCCGAAGAAATGGGTGGTTGGTTTGCATATCCTGGTAATGGCCAGACCAAAAATGCAACCGCAGTTAGTGGTAATGGTAATTCAGCAAGCAGCGCAACCGTTGTTTTTGGTGCAAAACGTCAAGAACTGGTGAAATAAAAGCTTCTGCCTGGCGGCCTTTTGCGCC

Table S2: Strain information for clinical isolates utilized for cross-reactivity analysis, including strain name, sex of patient the strain was recovered from, type of infection, Por type, and country of origin.

| Strain | Patient info | Isolation site | Por type | Country | Original Strain Name | Reference/Source |
| --- | --- | --- | --- | --- | --- | --- |
| 36343 | male | Invasive | porB1A | Canada |  | Public Health Ontario |
| 51748 | male | invasive | porB1A | Canada |  |  |
| 50412 | male | invasive | porB1A | Canada |  |  |
| 15Ngo01 | female | invasive | porB1B | Canada |  |  |
| 15Ngo08 | female | invasive | porB1A | Canada |  |  |
| 15Ngo15 | male | genital | porB1B | Canada |  |  |
| 17Ngo14 | female | genital | porB1B | Canada |  |  |
| 18Ngo10 | female | invasive | porB1A | Canada |  |  |
| 18Ngo16 | male | invasive | porB1A | Canada |  |  |
| 18Ngo18 | male | rectum | porB1A | Canada |  |  |
| N2082 | female | genital | PorB1A | Kenya | 4121 | Strains referenced in PMID: 10482498. PMID: 25605771 |
| N2067 | female | genital | unknown | Kenya | 4001 |  |
| N2078 | female | genital | PorB1A | Kenya | 4039 |  |
| N2061 | male | genital | unknown | Kenya | 2175 |  |
| N2071 | female | genital | PorB1B | Kenya | 4007 |  |
| N2066 | female | genital | unknown | Kenya | 2180 |  |
| N2080 | female | genital | PorB1B | Kenya | 4077 |  |
| N2075 | female | genital | unknown | Kenya | 4029 |  |
| N2068 | female | genital | PorB1A | Kenya | 4002 |  |
